# The cytidine deaminase APOBEC3C has unique sequence and genome feature preferences

**DOI:** 10.1101/2024.01.15.575754

**Authors:** Grant W. Brown

## Abstract

APOBEC proteins are cytidine deaminases that restrict the replication of viruses and transposable elements. Several members of the APOBEC3 family, APOBEC3A, APOBEC3B, and APOBEC3H-I, can access the nucleus and cause what is thought to be indiscriminate deamination of the genome, resulting in mutagenesis and genome instability. Although APOBEC3C is also present in the nucleus, its deamination target preferences are unknown. By expressing human *APOBEC3C* in a yeast model system, I have defined the APOBEC3C mutation signature, as well as the preferred genome features of APOBEC3C targets. The APOBEC3C mutation signature is distinct from those of the known cancer genome mutators APOBEC3A and APOBEC3B. APOBEC3C produces DNA strand-coordinated mutation clusters, and APOBEC3C mutations are enriched near the transcription start sites of active genes. Surprisingly, APOBEC3C lacks the bias for the lagging strand of DNA replication that is seen for APOBEC3A and APOBEC3B. The unique preferences of APOBEC3C constitute a mutation profile that will be useful in defining sites of APOBEC3C mutagenesis in human genomes.

---

The human *APOBEC3* family comprises seven paralogous genes that encode cytidine deaminases. APOBEC3 proteins target single-stranded DNA (ssDNA) and RNA to perform non-redundant functions in restricting the replication of viruses and the transposition of LINE elements (reviewed in (Koito and Ikeda 2013; Harris and Dudley 2015; Pecori *et al*. 2022)). More recently, several APOBEC3 proteins have been implicated in the mutagenesis of cancer genomes and in causing genome instability (reviewed in (Mertz *et al*. 2022)). Individual APOBEC3 proteins have diagnostic sequence and structural contexts for their preferred deamination sites, and the identification of these contexts has been instrumental in dissecting the contributions of individual APOBEC3 proteins to genome mutagenesis (Nik-Zainal *et al*. 2012; Roberts *et al*. 2012; Burns *et al*. 2013a; Alexandrov *et al*. 2013; Chan *et al*. 2015; Supek and Lehner 2017; Petljak *et al*. 2019, 2022; Buisson *et al*. 2019; Mas-Ponte and Supek 2020; Jakobsdottir *et al*. 2022; Sanchez *et al*. 2023).

Currently there is only a modest indication of the APOBEC3C deamination site consensus sequence. A preference for the T<u>C</u>A context (the underlined <u>C</u> being the deamination target) was noted in deamination of hepatitis B virus genomes by APOBEC3C (Khalfi *et al*. 2022), as was a preference for T or C at the -1 position relative to the deaminated deoxycytidine, and C at +1 (Chen *et al*. 2021). A preference for T or C at -1 was also noted in deamination assays of Moloney leukemia virus, as was some preference for T or A at -2 (Langlois *et al*. 2005). Purified APOBEC3C preferred T<u>C</u>A and T<u>C</u>G contexts *in vitro* (Ito *et al*. 2017). Analysis of 67 mutations accumulated in a yeast strain expressing APOBEC3C indicated a preference for T at -1 and at -2 (Taylor *et al*. 2013). By contrast, the deamination site consensus sequences for APOBEC3A and APOBEC3B are well understood (Chan *et al*. 2015; McDaniel *et al*. 2020; Hou *et al*. 2021; Petljak *et al*. 2022). Both prefer a T<u>C</u>W context (Taylor *et al*. 2013; Hoopes *et al*. 2016), as is typical of the APOBEC3 family (with the exception of A3G (Schumacher *et al*. 2005; Liu *et al*. 2023)), but differ in their preference at the -2 position (Chan *et al*. 2015). Together, these data suggest that the preferred deamination sites of APOBEC3A and APOBEC3B likely differ from that of APOBEC3C.

Prevailing models suggest that two factors contribute to deamination of the nuclear genome by APOBEC3 proteins: i) the intrinsic properties of each APOBEC3 protein, and ii) the features of the genome that could reveal substrates for deamination by APOBEC3s. Analyses in human cells, cancer genomes, and in humanized yeast models have established connections to genome features that are preferred sites of APOBEC3 action. Two themes predominate. First, there is an association with DNA replication and with active transcription (Kazanov *et al*. 2015; Morganella *et al*. 2016; Chervova *et al*. 2021). Second, there is a preference for regions presumed to have more single-stranded DNA: the lagging strand of DNA replication, the non-transcribed strand of transcription units, and DNA repair tracts (Haradhvala *et al*. 2016; Seplyarskiy *et al*. 2016; Hoopes *et al*. 2016; Morganella *et al*. 2016; Chervova *et al*. 2021; Chen *et al*. 2014; Mas-Ponte and Supek 2020; Sui *et al*. 2020). Whether these preferences are universal is unclear, and little is currently known about the genome features that might promote deamination by APOBEC3C.

Here I describe a humanized yeast model to define the deamination site consensus sequence and the preferred genome features for deamination by human APOBEC3C.

## Materials and Methods

### Yeast strains

Yeast strains (Table S1) used in this study were derived from BY4741 (Brachmann *et al*. 1998) with gene corrections to improve sporulation and mitochondrial function (Harvey *et al*. 2018). Strains were constructed using standard yeast genetic and molecular cloning methods and were cultured under standard conditions (Dunham *et al*. 2015). The *APOBEC3C* wild type and variant ORFs (C97S C100S and S188I) were assembled from PCR amplicons of codon-optimized synthetic DNAs containing the desired mutations. The gene expression transcription units were assembled from a hybrid pGPD14/15 promoter (Kotopka and Smolke 2020), the *APOBEC3C* open reading frame, sequences encoding the superFLAG epitope (Layton *et al*. 2019), and the tsynth27 transcription terminator (Curran *et al*. 2015), using Golden Gate assembly (HamediRad *et al*. 2019). Transcription units were integrated into the *LYP1* locus, targeted with a CRISPR/Cas9 double-strand break specified by an sgRNA sequence (5’-CATAATAACGTCCAATAAAT) cloned in the plasmid pUB1306 (constructed by Gavin Schlissel in Jasper Rine’s lab, and a kind gift from Elçin Ünal, UC Berkeley). APOBEC3 strains (wild type, C97S C100S, and S188I; strains GBY789, GBY793, and GBY805) were confirmed by PCR, by Sanger sequencing of PCR amplified gene loci, and by immunoblot analysis. *UNG1* was deleted by amplifying the *ung1Δ::kanMX* locus from genomic DNA purified from the *S. cerevisiae* genome deletion project *MAT***a** *ung1Δ::kanMX* strain, followed by transformation of the resulting PCR product into DHY214 to yield strain GBY732. The *ung1Δ* was crossed into the APOBEC3C strain followed by identification of an *ung1Δ::kanMX lyp1::A3C* segregant (GBY819). *RAD5* was deleted by amplifying the *rad5Δ::kanMX* locus from genomic DNA purified from the *S. cerevisiae* genome deletion project *MAT***a** *rad5Δ::kanMX* strain, followed by transformation of the resulting PCR product into DHY213 to yield strain GBY810.

### Immunoblot analysis

For validation of protein expression (Figure 1b), strains were grown to mid logarithmic phase and 5 OD_600_ units of cells were collected, diluted to a total volume of 10 ml in 10% trichloroacetic acid (TCA), incubated at room temperature for 15 minutes, then pelleted by centrifugation at 3000 rpm for 5 minutes. Fixed cells were washed with 1 ml of 1M Tris-HCl pH 6.8, harvested, resuspended in 50 µL of 5X Laemmli sample buffer (0.25M Tris-HCl pH 6.8, 0.5 M DTT, 10% SDS, 50% Glycerol, 0.5% bromphenol blue), and lysed by vortexing with 0.5mm glass beads for 10 minutes. An additional 50 µL 5x SDS-PAGE loading buffer was added to the sample and the extract was incubated at 95°C for 5 minutes and microfuged briefly prior to gel loading. Immunoblots were stained with 0.2% Ponceau S in 3% TCA, destained with TBST, and blocked with 5% non-fat dry milk in TBST. APOBEC3C tagged with the superFLAG epitope was detected by immunoblotting with an anti-FLAG mouse antibody (Sigma anti-FLAG M2 antibody F3165, 1:5000 dilution).

**Fig. 1.**
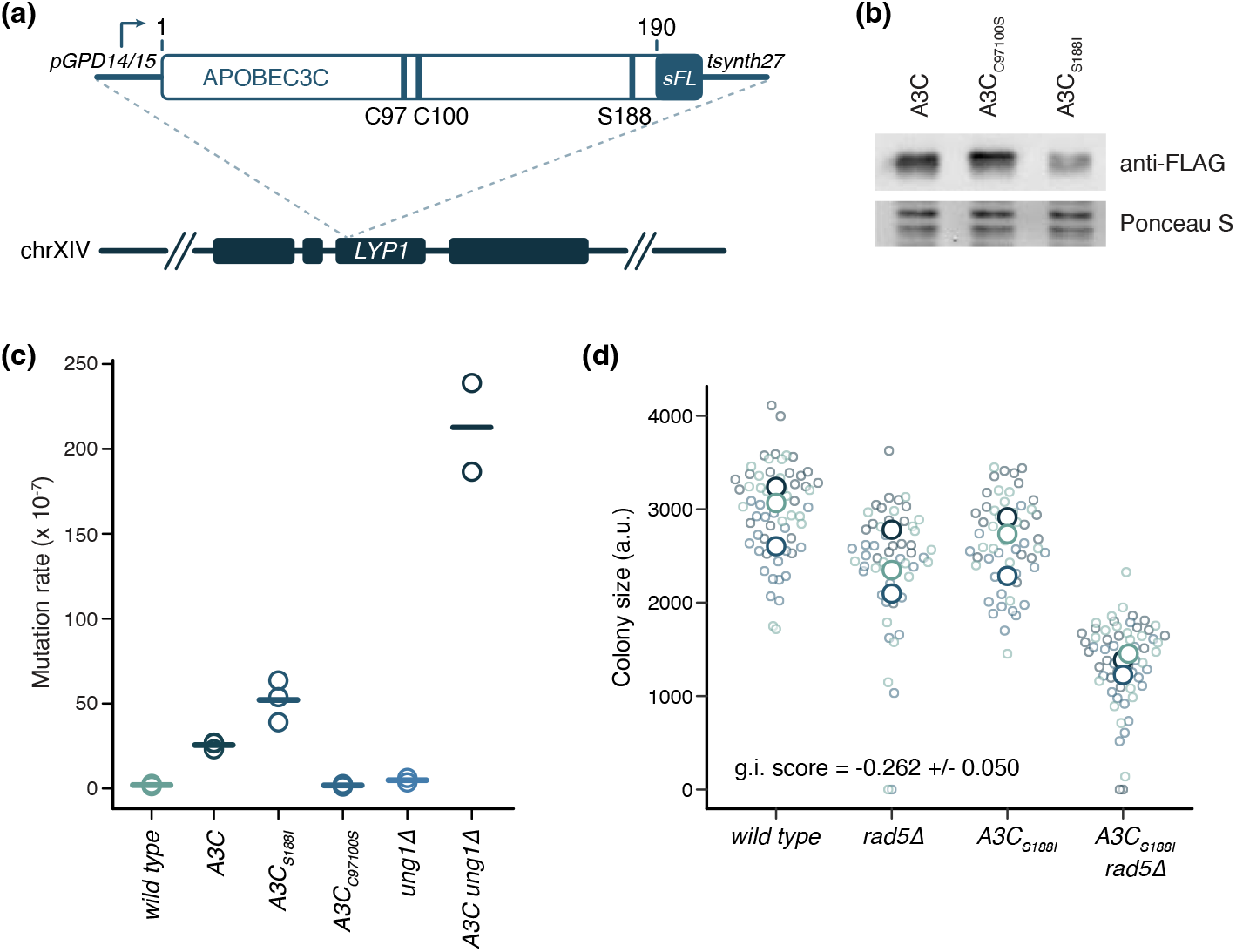
Expression of human *APOBEC3C* in yeast is mutagenic. a) Schematic diagram of the *APOBEC3C* expression construct. The promoter (pGPD14/15) and terminator (tsynth27) are indicated, as are the positions of the superFLAG (sFL) epitope tag, the catalytic-dead mutation (C97/C100), and the activating mutation (S188). The integration site in the *LYP1* gene is shown below. b) Immunoblot analysis of extracts of yeast strains expressing wild type (A3C), catalytic-dead (A3C_C97100S_), or hyperactive (A3C_S188I_) alleles of *APOBEC3C*. The blot was probed with anti-FLAG antibody to detect the A3C proteins. An image of the blot stained for total protein (Ponceau S) is shown as a loading control. c) Mutation rates at the *CAN1* locus were measured for yeast strains expressing the indicated *APOBEC3C* alleles. Where indicated (*ung1*Δ) the *UNG1* gene was deleted. d) Colony size in arbitrary units (a.u.) was measured for the indicated strains following dissection of tetrads from *rad5*Δ x *APOBEC3C*_*S188I*_crosses. Small circles show individual colony sizes (18 to 22 per genotype per replicate), and the large circles show the mean colony size for each of the 3 replicates. The genetic interaction (g.i.) score is indicated.

### Mutation rate assays

Mutation rates at *CAN1* were measured using a Luria-Delbrück fluctuation test (Lang 2018), with modifications. For each fluctuation test, saturated cultures were diluted 1:10,000 in 10 ml of SD-arginine, and 96 cultures (30 µl) were grown in a flat-bottom 96-well plate for 48 hours at 30°C without shaking. After incubation, 8 cultures were pooled, diluted with dH_2_O, and plated on fully supplemented SD to calculate N(t). For the remaining 88 cultures, 200 µL of SD-arginine containing 60 mg/l canavanine was added to each well. The plate was then sealed with an adhesive plate seal and incubated for 3 days at 30°C. Wells were scored for colony formation, and mutation rate was calculated according to the Poisson distribution (Table S2). Strains assayed were DHY213, GBY789, GBY793, GBY805, GBY818 and GBY819.

### Genetic interaction analysis

To assess genetic interactions between *APOBEC3C* alleles and *RAD5*, strains GBY789, GBY793, and GBY805 were each crossed to GBY810. Diploids of each were sporulated, and 20 tetrads of each were dissected on YPD. Plates were imaged after 48 hours at 30°C, and replica-plated to YPD + G418 to identify *rad5Δ* strains and to SD-lysine + S-(2-Aminoethyl)-L-cysteine hydrochloride to identify *APOBEC3C* strains. Colony sizes were measured from the plate images using Balony (Young and Loewen 2013), and raw values are listed in Table S3. The fitness of each genotype was calculated from the colony size values and expressed relative to the fitness of the wild type strain (Table S4). Genetic interaction scores were calculated as the deviation from expectation where expectation is the product of the single mutant fitness values (Table S4).

### Mutation accumulation

Strain GBY819, which expresses *APOBEC3C* in an *ung1Δ* background, was used for mutation accumulation. Eighteen independent lines were established from single colonies. Each line was streaked for single colonies on YPD and incubated at 30°C for 48 hours for one passage (approximately 25 generations). The single colony closest to the center of the plate was streaked for single colonies for each subsequent passage. After 15 passages (∼375 generations) or 20 passages (∼500 generations) the single colony closest to the center of the plate was inoculated into 2 ml YPD and incubated with shaking at 30°C for 48 hours. Genomic DNA was prepared from 0.5 ml of culture with the MasterPure Yeast DNA Purification kit (Biosearch Technologies) according to the manufacturer’s instructions. Genomic DNA concentrations were measured using Qubit dsDNA Broad Range reagent (ThermoFisher).

### Sequencing and variant calling

Sequencing libraries were prepared by tagmentation, using Nextera (Illumina) reagents scaled to small volume reactions (Vonesch *et al*. 2021). Libraries were pooled and subjected to paired-end sequencing with 75 bp reads on an Illumina NextSeq. De-multiplexed raw sequencing reads are available from the NCBI Sequencing Read Archive, BioProject ID PRJNA938955.

Variants were called for each mutation accumulation line using the Snippy tool in Galaxy (https://usegalaxy.org). Paired reads were concatenated and treated as single-end reads, the reference genome was saccCer3, and minimum read depth of 10 and minimum variant proportion of 0.9 were required to call a variant. Variant data were read into R. Any variant that occurred more than once was presumed to be parental and was removed. Complex variants and multiple nucleotide variants were manually deconvoluted into single nucleotide variants (SNVs). Variants mapping to the mitochondrial genome were removed. The resulting unique nuclear genome variants are tabulated in Table S5.

To make comparisons with APOBEC3A and APOBEC3B expressed in yeast, the data from Hoopes et al. were acquired (NCBI SRA BioProject ID PRJNA307256 and PRJNA307256), and variants were called with the VarScan Somatic tool in Galaxy, using saccCar3 as the reference genome, minimum coverage = 9, minimum supporting reads = 4, and minimum variant allele frequency = 0.4 (as these data were derived from diploid cells). Files (vcf) for SNVs were processed to retain heterozygous variants (TUMOR = 0/1) with FILTER = PASS, INFO = SOMATIC, and to remove variants that occurred more than once. SNVs mapping to the mitochondrial genome were removed. The resulting unique nuclear genome SNVs are tabulated in Tables S6 (APOBEC3A) and S7 (APOBEC3B).

SNVs resulting from deamination (C->T and G->A) were annotated and retained. SNVs in repetitive regions of the yeast genome were removed, using the definitions in (Sui *et al*. 2020).

### Mutation signatures

The three and seven nucleotide contexts of each deamination SNV were extracted using the mutSignatures package (Fantini *et al*. 2020) in R. Seven nucleotide contexts were analyzed with the web implementation of pLogo (O’Shea *et al*. 2013) (https://plogo.uconn.edu), using S288C_reference_sequence_ R64-3-1_20210421.fsa downloaded from SGD (https://www.yeastgenome.org) as the reference genome. Three nucleotide contexts were used to define the 96 trinucleotide mutation signatures for comparison to the COSMIC mutation signatures. Trinucleotide mutation signatures were corrected for trinucleotide frequency differences between the yeast and human genomes. The 96 trinucleotide signatures were plotted using the MutSignatures package (Fantini *et al*. 2020), and compared to the COSMIC signatures (COSMIC_v3.3.1_SBS_GRCh37 downloaded from https://cancer.sanger.ac.uk/signatures/downloads/ on March 3, 2023), using the MutationalPatterns package (Blokzijl *et al*. 2018).

### Mutation clustering

Clustering of deamination mutations was evaluated by calculating the distance between each pair of SNVs for each mutation accumulation line, yielding the intermutation distance for each pair. Each intermutation distance was annotated as strand-coordinated (both SNVs either C->T or G->A) or not. A previously-established cluster definition (Mas-Ponte and Supek 2020) of intermutation distance <500 bp was applied, with directly adjacent SNV pairs retained, and the likelihood of strand coordination was evaluated with a χ^2^ test. The expected number of deamination clusters was determined by randomly broadcasting the SNVs for each chromosome for each mutation accumulation line across the relevant chromosome, adjusting to the nearest C or G coordinate, before repeating the intermutation distance calculation and strand coordination evaluations. Randomization of SNVs was repeated 5 times to arrive at a median clustering expectation.

### Transcription bias analyses

To evaluate enrichment of APOBEC3 SNVs at different genomic features, potential APOBEC3A/B/C deamination targets were defined as the set of T<u>C</u> and <u>G</u>A dinucleotides in the sacCer3 reference genome and the coordinates of the C’s and G’s were extracted for use as the background reference set. The proportion of SNVs in genes and intergenic regions, and enrichment therein, was evaluated using the gene coordinates of the sacCer3 reference genome. To evaluate enrichment near start codons of genes, distance to nearest ATG and the strand of the nearest ATG was annotated for each SNV. Start codon proximal SNVs (+/- 500 bp) were collected in 50 bp bins, and the proportion of SNVs on the non-transcribed strand was calculated for each bin. A similar analysis was performed for tRNA genes, using the coordinates of the 275 tRNA genes in the reference genome, extracted using YeastMine (Balakrishnan *et al*. 2012) (https://yeastmine.yeastgenome.org; accessed on 1/24/2023). Analysis of proximity to highly expressed genes utilized the gene expression table in Supplementary Dataset S3-4 from (Sui *et al*. 2020), from which the highest and lowest 5% of expressed genes were extracted after removing genes whose expression was not detected.

### Replication bias analyses

Replication timing and strand bias was evaluated by first calculating the distance between each SNV and the nearest early-firing DNA replication origin (Balint *et al*. 2015) or the nearest efficient origin (efficiency > 0.7) (McGuffee *et al*. 2013). As the results were highly similar, calculations from the (Balint *et al*. 2015) dataset were subjected to further analyses. SNVs within +/- 7 kb of an early origin were collected in 700 bp bins for visualization, and enrichment for early origin proximity was evaluated by χ^2^ tests. Replication strand bias was evaluated using early-firing origin annotations (Balint *et al*. 2015) without normalization of inter-origin distances, as the certainty of replication strand (leading or lagging) diminishes with the actual distance from an efficient early origin. Distances-to-origin were collected in 5 kb bins for 50 kb flanking each early origin and the proportion of C->T SNVs was calculated for each bin. C->T SNVs to the right of an origin and G->A SNVs to the left of an origin are defined as lagging strand template deaminations. As for the transcription bias analyses, APOBEC3 targets were defined as TC or GA dinucleotides.

### Computational Analyses, Software, and Data Availability

Statistical analysis, data manipulation, and data visualization were performed using the R Statistical Software (R Core Team 2021) in RStudio. Figures were assembled using Adobe Illustrator v28.1 using the Lost in Translation colour palette (https://www.instagram.com/filmandcolor/). Strains and plasmids are available upon request. Raw sequencing reads from the *APOBEC3C* mutation accumulation experiment are available from the NCBI Sequencing Read Archive, BioProject ID PRJNA938955. The author affirms that all data necessary for confirming the conclusions of the article are present within the article, figures, tables, and supplementary materials.

## Results

### A humanized yeast platform to study APOBEC3C-mediated mutagenesis

To rigorously define the preferred sites of deamination by APOBEC3C, I took advantage of the approach used successfully for APOBEC3G (Schumacher *et al*. 2005), APOBEC3A, and APOBEC3B (Hoopes *et al*. 2016), and expressed APOBEC3C constitutively in the budding yeast *S. cerevisiae*. I flanked a yeast codon-optimized *APOBEC3C* open reading frame with a strong constitutive artificial promoter (Kotopka and Smolke 2020) and a synthetic terminator (Curran *et al*. 2015) (Fig. 1a). The *APOBEC3C* transcription unit was then integrated in single copy in the *LYP1* gene to provide optimal stability. In addition to the wild type *APOBEC3C* gene, I also expressed a catalytic-dead variant encoding C97S and C100S (Horn *et al*. 2014), and a naturally-occurring hyperactive allele encoding S188I (Duggal *et al*. 2013; Wittkopp *et al*. 2016).

Expression of the three APOBEC3C proteins was confirmed by immunoblot analysis (Fig. 1b). Wild type and catalytic-dead proteins were expressed at similar levels, while the hyperactive allele was expressed at lower levels. To test whether APOBEC3C expressed in yeast was mutagenic, I performed fluctuation analyses to measure the forward mutation rate at the *CAN1* locus (Fig. 1c). As anticipated, APOBEC3C expression increased the mutation rate, from 2 × 10^−7^ to 26 × 10^−7^. Expression of the hyperactive S188I variant further increased mutation rate to 52 × 10^−7^. Expression of the catalytic-dead C97S/C100S mutant did not increase the mutation rate (1.8 × 10^−7^). To test whether the mutagenesis by APOBEC3C was the result of cytidine deamination a deletion of the *UNG1* gene, which encodes the yeast uracil DNA glycosylase, was introduced into the APOBEC3C expression strain. Deletion of *UNG1* resulted in a large increase in mutation rate, to 213 × 10^−7^, as would be expected when removal of uracils by base excision repair is prevented following cytidine deamination. As a final measure of APOBEC3C function *in vivo* in the humanized yeast system, I introduced a deletion of the *RAD5* gene into the APOBEC3C expression strains and measured cell fitness (Fig. 1d, Supplementary Fig. 1a and 1b). Deletion of *RAD5* results in accumulation of ssDNA in the nucleus (Gallo *et al*. 2019), and results in fitness defects when combined with expression of APOBEC3A or APOBEC3B (Hoopes *et al*. 2016). Although *APOBEC3C* expression in *rad5Δ* resulted in only a modest fitness defect (Supplementary Fig. 1a), expression of the hyperactive S188I variant resulted in a fitness defect that was much greater than expectation (Fig. 1d; multiplicative model, comparing to *rad5Δ* and *APOBEC3C* expression alone). I conclude that APOBEC3C expressed in yeast is deaminating nuclear DNA, likely ssDNA, to produce mutations.

### Mutation accumulation during APOBEC3C expression

To analyse the sites deaminated by APOBEC3C *in vivo* I established 18 clonal lineages originating from the same *ung1Δ APOBEC3C* expression strain, propagated them independently for 375 to 500 generations to allow mutations to accumulate, and imposed a single cell bottle-neck every ∼25 generations to minimize the effects of genetic drift (Zhu *et al*. 2014). Genomic DNA from each strain was deep sequenced, and a total of 6771 unique variants were identified (Fig. 2a). After removing variants that mapped to repetitive regions of the genome, there were 6692 variants, the vast majority of which (6685) were single-nucleotide variants (SNVs) resulting from deamination. The average number of deamination SNVs per mutation accumulation line was 370 (Fig. 2b). I also analysed the mutation accumulation data reported for APOBEC3A and APOBEC3B expressed in yeast (Hoopes *et al*. 2016) to facilitate comparative analyses, identifying 1421 and 1643 deamination SNVs, respectively (Fig. 2b). The average mutation rate across the 18 APOBEC3C mutation accumulation lines was 0.73 × 10^−7^ per base per generation (Fig. 2c), about 400-fold the wild type rate (Zhu *et al*. 2014), and about half that observed upon APOBEC3B expression in yeast (Sui *et al*. 2020).

**Fig. 2.**
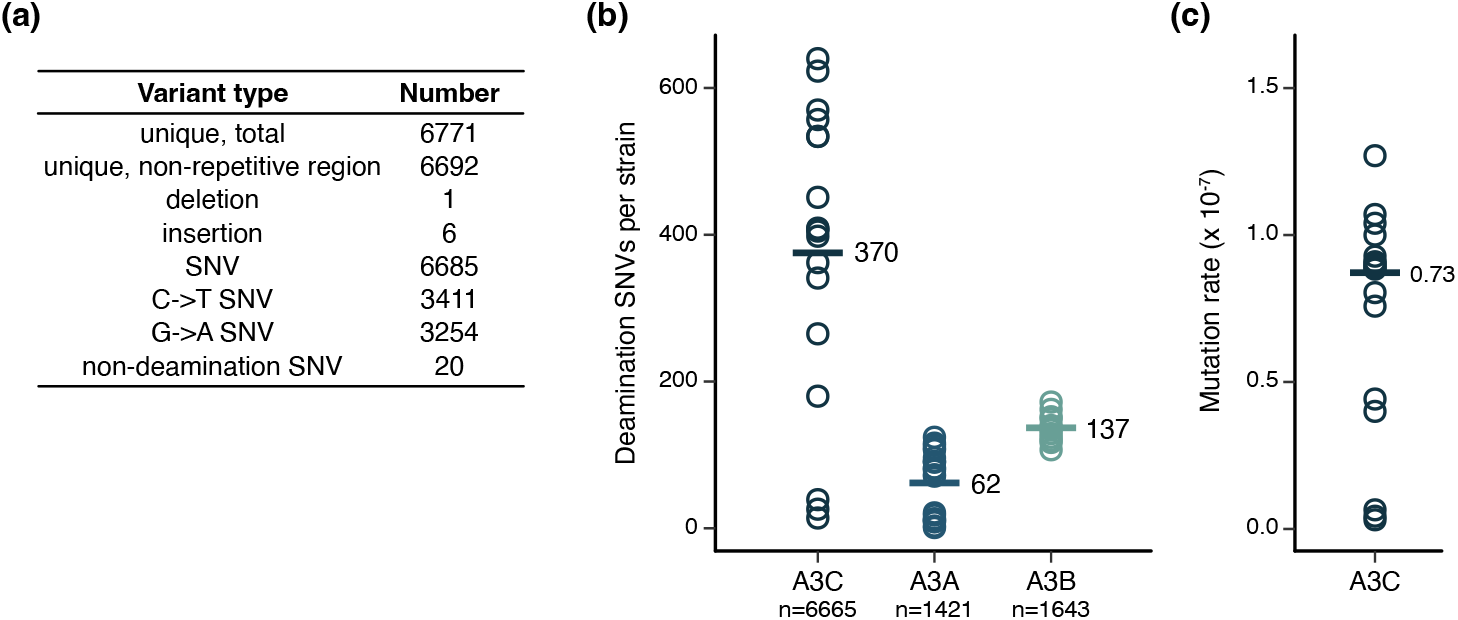
Mutation accumulation in yeast cells expressing *APOBEC3C*. a) Summary of the variant types in the 18 mutation accumulation lines. The number of single-nucleotide variants (SNV) and the type of SNV (C->T or G->A) are indicated. b) The number of SNVs due to deamination is plotted for each mutation accumulation line for yeast expressing *APOBEC3C, 3A*, or *3B*. Horizontal bars indicate the means, and the total number of deamination SNVs for each mutation accumulation experiment is indicated. *APOBEC3A* and *3B* data are from (Hoopes *et al*. 2016). c) The mutation rate is plotted for each of the 18 *APOBEC3C* mutation accumulation lines. The horizontal bar indicates the mean.

### The APOBEC3C mutation signature differs from other APOBEC3 proteins

Using the APOBEC3C mutation accumulation data, along with the data reported by Hoopes *et al* for APOBEC3A and APOBEC3B (Hoopes *et al*. 2016), I extracted the genome contexts for each unique deamination and identified over- and under-represented bases at positions flanking the deaminated deoxycytidine (Fig. 3a). As is typical of APOBEC3 proteins (with the exception of APOBEC3G) APOBEC3C showed a strong preference for T at the -1 position. Both A and G were under-represented at -1, while C was neutral. At -2, the preference for T indicated by Taylor *et al* (Taylor *et al*. 2013) is apparent, as is a very strong under-representation of C and G. The motif derived for APOBEC3C extends 3 nucleotides upstream and downstream of the target C, with under-representation of G and C apparent at -3, +1, +2, and +3. Examination of the motif for APOBEC3C revealed clear differences from APOBEC3A and APOBEC3B (Fig. 3a). The motifs that I derived for APOBEC3A and APOBEC3B were similar to the TCW trinucleotide motif reported in the original analysis of the datasets (Hoopes *et al*. 2016), and were similar to the extended five nucleotide motifs reported by Chan *et al* (Chan *et al*. 2015) and to the preferences at -2 reported by Taylor *et al* (Taylor *et al*. 2013). The APOBEC3C motif differs most at the -2 position, with an over-representation of T that is not observed for APOBEC3B and that is stronger than APOBEC3A. APOBEC3C also shows an under-representation of C at -2 that is not apparent for either APOBEC3A or APOBEC3B. Additionally, C is neutral at -1 in the APOBEC3C motif, whereas it is under-represented in the APOBEC3A and APOBEC3B motifs. The differences in deamination motifs between APOBEC3C, APOBEC3A, and APOBEC3B are also evident in the conventional triplet motif representation of their mutation signatures (Fig. 3b and Supplementary Fig. 2), where the high fractions of T<u>C</u>A and T<u>C</u>T variants seen for APOBEC3A and APOBEC3B are substantially diminished for APOBEC3C. The C<u>C</u>A, C<u>C</u>T, and C<u>C</u>C variants each approach 10% for APOBEC3C (9.4%, 7.7%, and 9.8%), while making up fewer than 3% of APOBEC3A variants, and fewer than 4% of APOBEC3B variants.

**Fig. 3.**
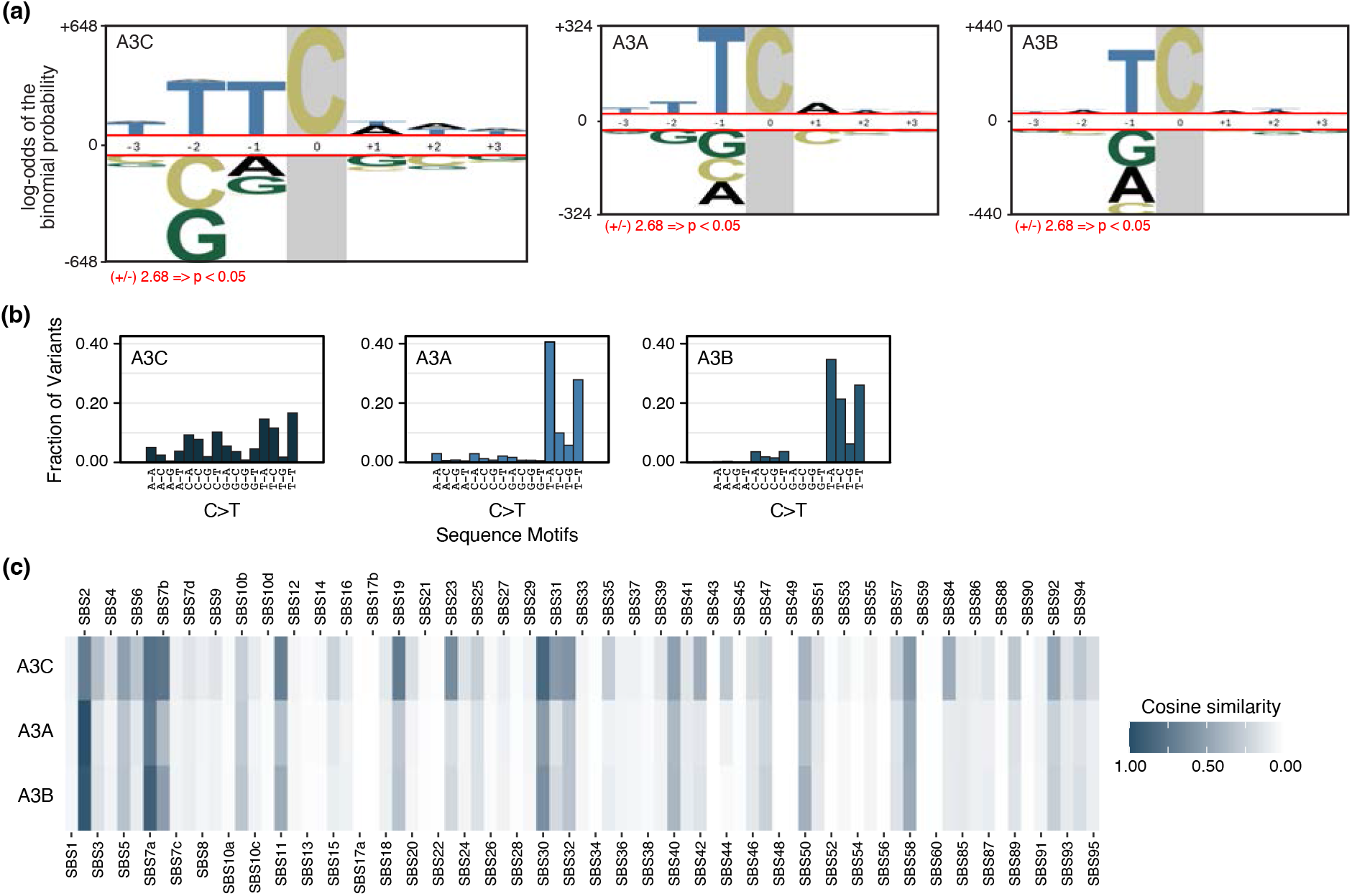
The mutation signature of APOBEC3C is distinct from that of APOBEC3A or 3B. a) A probability logo visualization of the APOBEC3C, 3A, and 3B deamination sequence motifs. Nucleotides are scaled relative to their log-odds binomial probabilities, with over-represented nucleotides above the x-axis. Red lines indicate the Bonferroni-corrected support threshold of p < 0.05. b) The tri-nucleotide mutational signatures are plotted for APOBEC3C, 3A, and 3B. Mutation types are grouped with only the 16 SNV types centered on C to T mutations shown. c) Heatmap showing the cosine similarity of the mutational signatures in (b) with the COSMIC single base substitution (SBS) signatures.

Having defined the mutational signature for APOBEC3C, I compared the trinucleotide frequencies in the APOBEC3C signature to the COSMIC single-base substitution (SBS) mutational signatures (Fig. 3c). As anticipated, APOBEC3A and APOBEC3B both had the highest similarity to SBS2, a signature attributed to APOBEC3A and APOBEC3B activities (Nik-Zainal *et al*. 2012; Chan *et al*. 2015; Petljak *et al*. 2022). By contrast, the APOBEC3C mutational signature is most similar to SBS30, a signature attributed to base excision repair deficiency (Drost *et al*. 2017), and had a higher similarity to SBS11, SBS19, and SBS23 than either APOBEC3A or APOBEC3B. Similarities to SBS13 were absent as all mutation accumulation strains were deficient in base excision repair. Together, these data indicate that APOBEC3C generates a distinct mutational profile.

### APOBEC3C mutations are found in clusters

APOBEC mutations in cancer genomes are associated with interesting patterns, where regional clustering of SNVs is observed, often with mutations occurring on the same DNA strand (Nik-Zainal *et al*. 2012). Two clustering patterns have been described: kataegis, where local strand-coordinated hypermutation events of tens of mutations occur over a span of kilobases (Nik-Zainal *et al*. 2012; Taylor *et al*. 2013), and omikli, where mutation clusters occur in pairs or triplets with a shorter inter-mutation distance (Mas-Ponte and Supek 2020). I measured the distances between APOBEC3C SNVs for each mutation accumulation line to assess the extent of clustering (Fig. 4a). Kataegis is expected to be rare in the APOBEC3C dataset as yeast models indicate that kataegis requires the action of uracil DNA glycosylase (Taylor *et al*. 2013), although kataegis in human cells does not require uracil DNA glycosylase (Petljak *et al*. 2022). The APOBEC3C yeast strains used here carry complete deletion of the uracil DNA glycosylase gene *UNG1*. Indeed, there were no mutation clusters that met a reasonable definition of kataegis (≥ 5 consecutive mutations with intermutation distances of ≤1 kb (Alexandrov *et al*. 2013; Petljak *et al*. 2019; Mas-Ponte and Supek 2020)). Clustered APOBEC3C SNV pairs, defined as having intermutation distances ≤500 bp (Supek and Lehner 2017; Mas-Ponte and Supek 2020), were more likely to occur on the same DNA strand than were non-clustered mutations (χ^2^ test, p=1 × 10^−10^). Strand coordination was particularly likely for SNV pairs with the shortest intermutation distances (Fig. 4b). All strand-coordinated mutation clusters caused by APOBEC3C, of which there were 308, consisted of pairs or triplets, reminiscent of omikli (Mas-Ponte and Supek 2020). APOBEC3C caused strand-coordinated clustered mutations at a high frequency (4.8% of SNV pairs were clustered, expectation=1.0%). By contrast, APOBEC3A and APOBEC3B caused clustered mutations at lower frequencies (1.1% and 1.3%, respectively, Fig. 4c and 4d). APOBEC3B clustered mutations were more likely to be strand coordinated (χ^2^ test, p=0.009), whereas APOBEC3A clustered mutations were not (χ^2^ test, p=0.5). Absence of strand coordination of APOBEC3A clustered mutations is consistent with the lack of processivity of APOBEC3A *in vitro* (Love *et al*. 2012), and the presence of strand coordination for APOBEC3B and APOBEC3C is consistent with the processive nature of both deaminases (Adolph *et al*. 2017a; b).

**Fig. 4.**
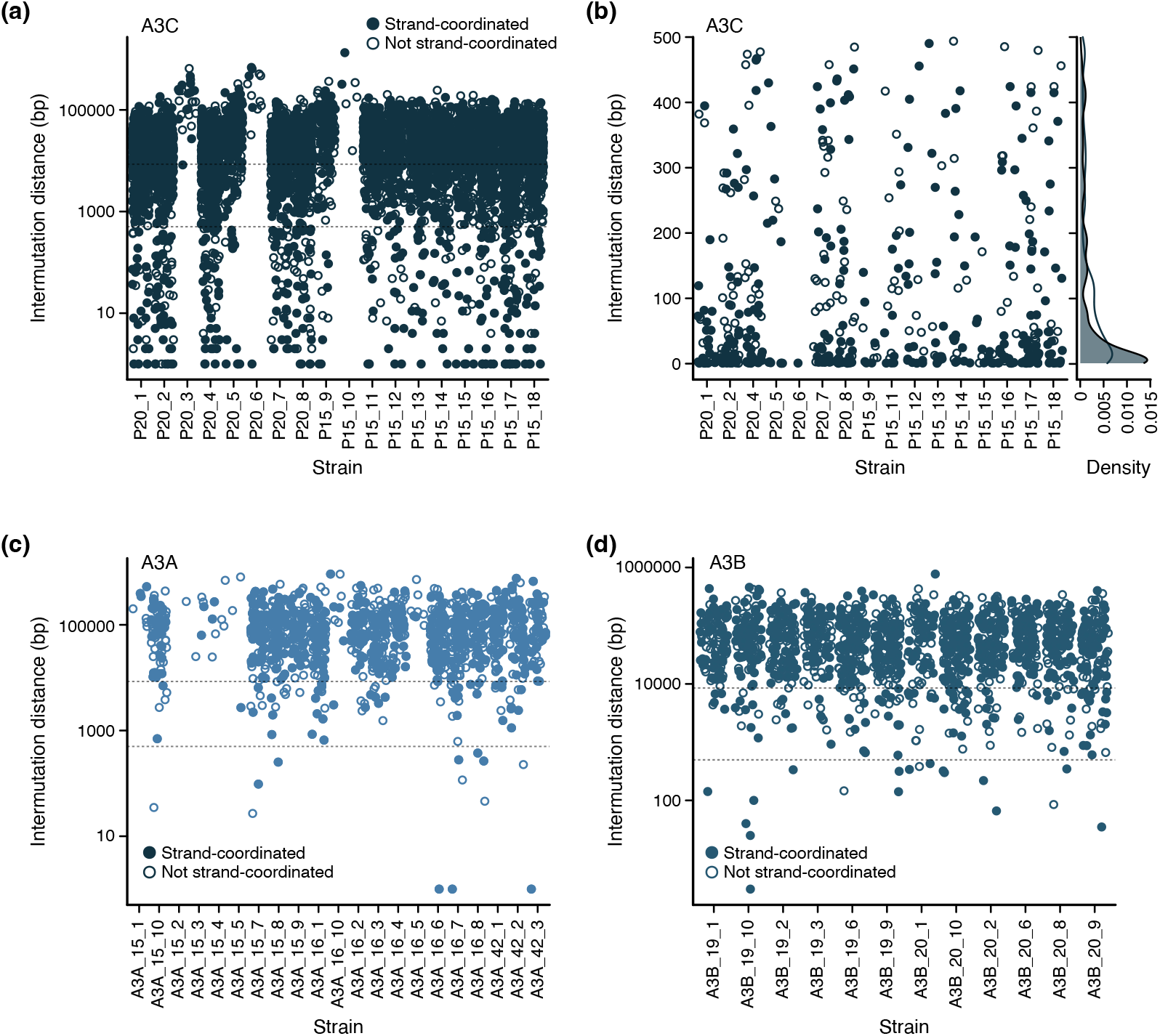
APOBEC3C makes clustered strand-coordinated mutations. a) The intermutation distance is plotted for each pair of APOBEC3C SNVs for each mutation accumulation line. Mutation pairs on the same DNA strand are indicated by closed circles. The horizontal dotted lines indicate intermutation distances of 8500 bp and 500 bp as cutoffs for proximal and clustered mutations, respectively. b) The intermutation distance is plotted for each pair of APOBEC3C SNVs for each mutation accumulation line, for intermutation distances less than 500 bp. Mutation pairs on the same DNA strand are indicated by closed circles. The density distributions of the strand-coordinated intermutation distances (shaded histogram) and the non-strand-coordinated intermutation distances (open histogram) are plotted on the right. c) The intermutation distance is plotted for each pair of APOBEC3A SNVs for each mutation accumulation line. Details as in (a). d) The intermutation distance is plotted for each pair of APOBEC3B SNVs for each mutation accumulation line. Details as in (a).

### APOBEC3C targets both strands of RNA polymerase II and tRNA genes

Transcription can expose ssDNA substrates for ectopic action by cytidine deaminases in humanized yeast models (Taylor *et al*. 2014; Lada *et al*. 2015; Hoopes *et al*. 2016). There is some controversy as to whether exposure of ssDNA by transcription is also important for APOBEC3 action in its native context, as assessed by cancer genome sequencing (Kazanov *et al*. 2015; Mas-Ponte and Supek 2020; Chervova *et al*. 2021). Given the compact nature of the yeast genome, intergenic regions tend to comprise promoter elements and transcription start sites (TSS), with the latter being in close proximity to the start codon of the open reading frame (median ∼40 bp (Ronsmans *et al*. 2019; Lu and Lin 2019)). I quantified deaminations across the start codon of open reading frames (± 500 bp) for APOBEC3C, APOBEC3A, and APOBEC3B (Fig. 5a). Both APOBEC3C and APOBEC3A showed deamination peaks in the TSS (−100 to -50 bp) and in the 5’ region of the open reading frame (+50 to +100 bp). By contrast, APOBEC3B deaminations were not enriched in regions proximal to start codons (Fig. 5a and 5b). I examined the strand biases of APOBEC3C and APOBEC3A deaminations that were proximal to start codons (Fig. 5c) and observed an interesting difference. APOBEC3A deaminations were biased to the non-transcribed strand, as previously noted (Hoopes *et al*. 2016). By contrast, APOBEC3C deaminations were biased to the transcribed strand, particularly within the first ∼100 bp of the open reading frame (Fig. 5c).

**Fig. 5.**
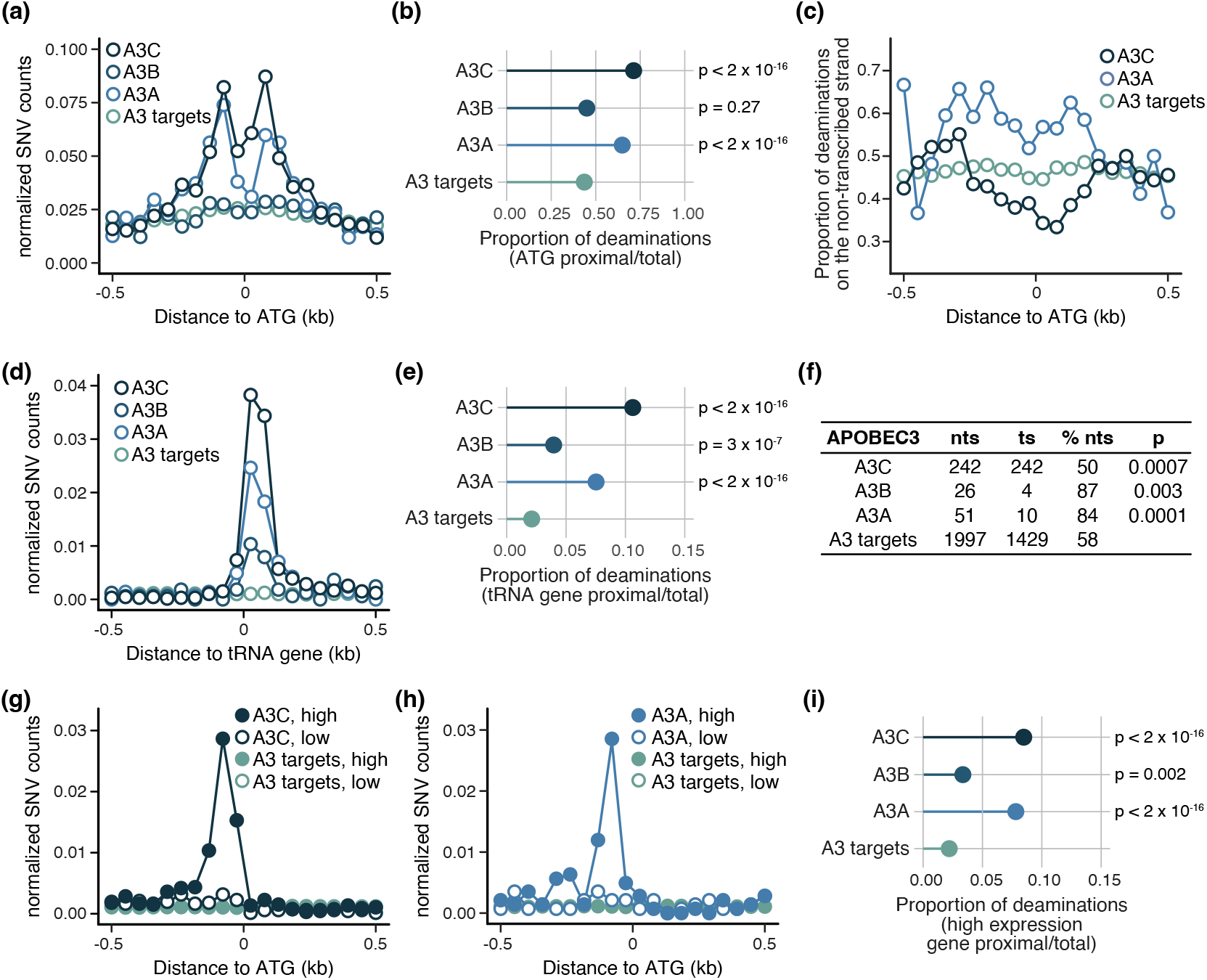
APOBEC3C targets the 5’ ends of pol II and tRNA genes. a) The proportion of SNVs in 50 bp bins surrounding pol II gene start codons is plotted for each APOBEC3. Putative APOBEC3 targets (T<u>C</u>) are plotted as a control. b) The proportion of deaminations +/- 500 bp of pol II gene start codons is plotted for APOBEC3C, 3B, and 3A. Putative APOBEC3 targets (T<u>C</u>) are plotted as a control. Statistical support for enrichments proximal to ATGs relative to APOBEC3 targets was assessed with chi-squared tests. c) The proportion of deaminations on the nontranscribed strand in 50 bp bins surrounding pol II gene start codons is plotted for each APOBEC3. Putative APOBEC3 targets (T<u>C</u>) are plotted as a control. d) The proportion of SNVs in 50 bp bins surrounding tRNA genes is plotted for each APOBEC3. Putative APOBEC3 targets (T<u>C</u>) are plotted as a control. e) The proportion of deaminations +/- 500 bp of tRNA genes is plotted for APOBEC3C, 3B, and 3A. Putative APOBEC3 targets (T<u>C</u>) are plotted as a control. Statistical support for enrichments proximal to tRNA genes relative to APOBEC3 targets was assessed with chi-squared tests. f) Deaminations by APOBEC3C, 3A, and 3B on the nontranscribed and transcribed strand +/- 500 bp of tRNA genes is tabulated. Statistical support for enrichments relative to APOBEC3 targets was assessed with chi-squared tests. g) The proportion of SNVs in 50 bp bins surrounding the highest 5% (closed circles) and the lowest 5% of expressed genes is plotted for APOBEC3C. Putative APOBEC3 targets (T<u>C</u>) are plotted as a control. h) The proportion of SNVs in 50 bp bins surrounding the highest 5% (closed circles) and the lowest 5% of expressed genes is plotted for APOBEC3A. Putative APOBEC3 targets (T<u>C</u>) are plotted as a control. i) The proportion of deaminations +/- 500 bp of the most highly expressed genes is plotted for APOBEC3C, 3B, and 3A. Putative APOBEC3 targets (T<u>C</u>) are plotted as a control. Statistical support for enrichments proximal to highly expressed genes relative to APOBEC3 targets was assessed with chi-squared tests.

APOBEC3B deaminations are enriched proximal to tRNA genes on the non-transcribed strand (Saini *et al*. 2017; Sui *et al*. 2020). I examined the propensity for APOBEC3C, APOBEC3A, and APOBEC3B to catalyze deaminations near tRNA genes (Fig. 5d). All three deaminases caused peaks of deamination in the first 75 bp of tRNA genes, as well as strong enrichments for tRNA gene-proximal deamination (Fig. 5e). APOBEC3B had the previously-described bias for the non-transcribed strand (Saini *et al*. 2017; Sui *et al*. 2020), as did APOBEC3A (Fig. 5f). By contrast, APOBEC3C deaminations occurred in identical numbers on the transcribed and non-transcribed strands, resulting in a small bias for the transcribed strand relative to that of APOBEC3 targets. I infer that APOBEC3C lacks the strong bias for the non-transcribed strand that is seen for APOBEC3A and APOBEC3B.

Since tRNA genes are highly-expressed (Warner 1999), I asked whether highly-expressed RNA pol II genes were particularly susceptible to deamination by APOBEC3C (Fig. 5g and 5i). APOBEC3C deaminations showed a clear peak at the TSS of highly-expressed pol II genes (the highest 5% of genes (Sui *et al*. 2020); Fig. 5g), as did APOBEC3A deaminations (Fig. 5h). Deaminations by APOBEC3C and APOBEC3A were enriched by 3.9- and 3.5-fold near highly expressed genes (Fig. 5i), whereas APOBEC3B deaminations showed a more modest 1.5-fold enrichment (Fig. 5i and Supplementary Fig. 3a). None of the APOBEC3 deaminations were enriched near the lowest expressed genes (Supplementary Fig. 3b).

### APOBEC3C lacks a DNA replication strand bias

APOBEC3 mutagenesis has been associated with early-replicating regions in human cancer genomes (Kazanov *et al*. 2015; Chervova *et al*. 2021). However, the association with replication timing might differ for clustered versus scattered mutations (Seplyarskiy *et al*. 2016) and is not evident in all analyses (Morganella *et al*. 2016; Buisson *et al*. 2019). Given the extensive replication origin annotations for the yeast genome (Siow *et al*. 2012), as well as precise origin timing and efficiency datasets (Siow *et al*. 2012; McGuffee *et al*. 2013), humanized yeast systems are ideal for analyzing relationships between DNA replication and APOBEC3 mutagenesis. I first asked if APOBEC3 deaminations were enriched in regions proximal to early-firing DNA replication origins (Fig. 6a, 6b, 6c, and Supplementary Fig. 4a), since the regions closest to early-firing origins will be the earliest to replicate. Both APOBEC3C and APOBEC3A showed a modest enrichment of 1.2-fold in regions +/- 7 kb of an early-firing origin, relative to potential APOBEC3 targets (Fig. 6c). APOBEC3B did not show a statistically supported enrichment proximal to early-firing origins (Fig. 6c).

**Fig. 6.**
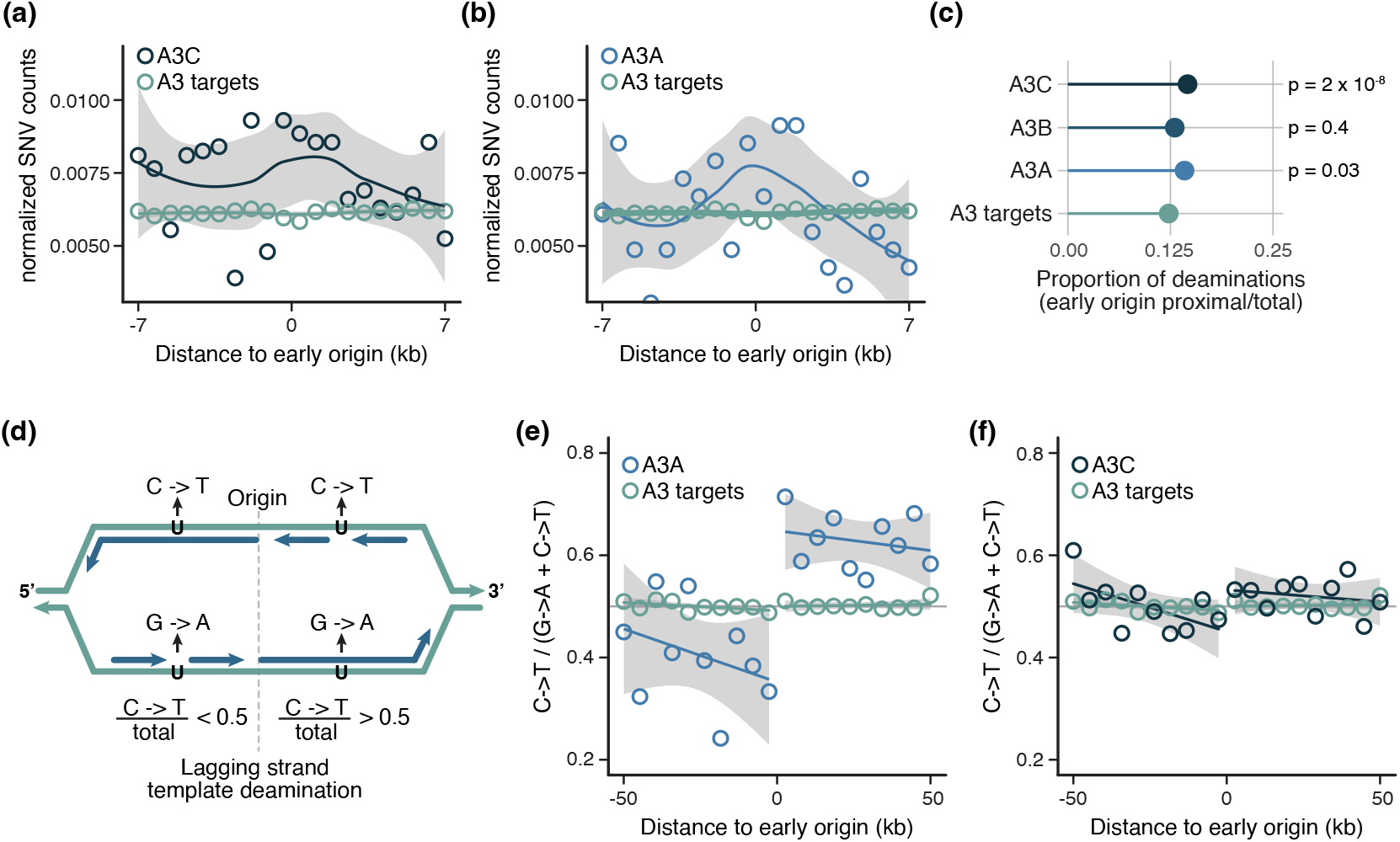
APOBEC3C lacks a DNA replication strand preference. a) The proportion of SNVs in 70 bp bins surrounding early-firing DNA replication origins is plotted for APOBEC3C. Putative APOBEC3 targets (T<u>C</u>) are plotted as a control. The solid line is the loess smoothed conditional mean with the shaded regions showing the standard error with a 0.95 confidence interval. b) The proportion of SNVs in 70 bp bins surrounding early-firing DNA replication origins is plotted for APOBEC3A. Details as in (a). c) The proportion of deaminations in regions proximal to early-firing origins is plotted for APOBEC3C, 3B, and 3A. Putative APOBEC3 targets (T<u>C</u>) are plotted as a control. Statistical support for enrichments in origin-proximal regions relative to APOBEC3 targets was assessed with chi-squared tests. d) Schematic diagram depicting the change in replication strand bias on opposite sides of a replication origin. To the left of the origin, a deamination on the top strand results in a C->T transition in the genome sequencing data, whereas a deamination on the bottom strand results in a G->A transition since the genome sequences are mapped onto the top strand. To the left of the origin, the bottom strand is the lagging strand template and so a preference for lagging strand template deamination manifests as a reduced proportion of C->T transitions. To the right of the origin, the opposite is true because now the top strand is the lagging strand template. e) The proportion of C->T SNVs in 5 kbp bins surrounding early-firing DNA replication origins is plotted for APOBEC3A. Putative APOBEC3 targets (T<u>C</u>) are plotted as a control. The solid line is the loess smoothed conditional mean with the shaded regions showing the standard error with a 0.95 confidence interval. f) The proportion of C->T SNVs in 5 kbp bins surrounding early-firing DNA replication origins is plotted for APOBEC3C. Details as in (e).

Data from cancer genomes (Haradhvala *et al*. 2016; Seplyarskiy *et al*. 2016) and from humanized yeast systems (Hoopes *et al*. 2016; Sui *et al*. 2020) indicate that APOBEC3A and APOBEC3B deaminations are biased toward the lagging strand DNA replication template. I asked whether APOBEC3C showed a similar bias (Fig. 6d, 6e, 6f, and Supplementary Fig. 4b). I considered genome regions located +/- 50 kb from early-firing origins, as the direction of replication fork movement, and therefore the assignment of leading versus lagging strand, is most certain in proximity to early origins. Since SNVs are annotated relative to the genome reference strand, replication strand bias can be visualized as a change in the proportion of C->T transitions as the direction of fork movement changes from right-to-left to left-to-right at replication origins (Fig. 6d). Lagging strand template bias results in a higher proportion of C->T to the right of origins and a lower proportion to the left. As previously described, both APOBEC3A (Fig. 6e) and APOBEC3B (Supplementary Fig. 4b) display a clear preference to deaminate the lagging strand template, and the bias is stronger closer to early-firing replication origins. Surprisingly, APOBEC3C does not have a substantial bias and deaminates the leading and lagging strand templates approximately equally (Fig. 6f).

## Discussion

Through my analysis of APOBEC3C deamination mutations in a yeast model, several interesting patterns of APOBEC3C activity have emerged that clearly distinguish APOBEC3C from APOBEC3A and APOBEC3B. The APOBEC3C mutation signature is distinct, with an over-representation of T at -2 and decreased bias against C at -1, relative to the APOBEC3A or APOBEC3B signatures. APOBEC3C produces strand-coordinated mutation clusters that are consistent with processive deamination. The absence of replication and transcription strand biases seen with other APOBEC3s further distinguishes the properties of APOBEC3C.

The unique mutation signature of A3C has some interesting implications. Explorations of the role of APOBEC3 proteins in mutating cancer genomes have largely focused on APOBEC3A and 3B (Burns *et al*. 2013a; b; Taylor *et al*. 2013; Alexandrov *et al*. 2013; Chan *et al*. 2015; Petljak *et al*. 2021, 2022; Isozaki *et al*. 2023; Caswell *et al*. 2023), with some attention paid to APOBEC3H haplotype I (Starrett *et al*. 2016), APOBEC3G (Liu *et al*. 2023), and APOBEC3C (Qian *et al*. 2022). Indeed, recent analyses indicate that APOBEC3A is the predominant mutator of cancer genomes, with a lesser role for APOBEC3B (Petljak *et al*. 2022), although the relative contributions likely vary in different contexts (Carpenter *et al*. 2023). Removing both APOBEC3A and APOBEC3B does not eliminate mutagenesis (Petljak *et al*. 2022), indicating that additional APOBEC3 proteins could also cause mutations in tumour cells. A common feature of cancer genome analyses with respect to APOBEC3 is the use of the T<u>C</u>N or T<u>C</u>W context, which manifests as mutation signatures SBS2 and SBS13, to identify mutations with a high likelihood of being caused by APOBEC3 activity (Green and Weitzman 2019; Petljak and Maciejowski 2020). However, while the T<u>C</u>N context captures 86% of APOBEC3A and 91% of APOBEC3B mutations analysed here, T<u>C</u>N captures only 50% of the APOBEC3C mutations. Thus, it is possible that the mutagenic influence of APOBEC3C has been underestimated in current cancer genome analyses. It will be of great interest to mine cancer genome data for evidence of APOBEC3C mutation, using the mutation signature derived in the humanized yeast model.

The features that dictate mutagenesis by APOBEC3 deaminases are subjects of great interest that has focused on two realms. The first includes explorations of contexts that could generate or reveal ssDNA substrates for APOBEC3 mutagenesis and the second examines intrinsic features of the APOBEC3 enzymes. One context that is likely to be critically important in revealing ssDNA substrates for APOBEC3 deaminases is DNA replication stress. The biases to early replicating regions (Fig. 6a, b, and c), tRNA genes (Fig. 5d and e), and highly expressed pol II genes (Fig. 5g and h) that are evident in the humanized yeast models are typically of a modest effect size, consistent with the presence of ssDNA being at native levels in these contexts. Introducing mutations known to produce abnormal levels of ssDNA, such as deleting *RAD5* (Gallo *et al*. 2019), increases mutagenesis by APOBEC3 proteins (Figure 1d and (Hoopes *et al*. 2016)), as do mutations that cause DNA replication stress (Mertz *et al*. 2023). Similarly, DNA replication stress induced in human cells by gemcitabine results in a dramatic increase in cytidine deamination [Ubhi 2024, in press]. The biases toward early replicating regions and the lagging strand that are evident in cancer genomes (Green *et al*. 2016; Haradhvala *et al*. 2016; Seplyarskiy *et al*. 2016; Morganella *et al*. 2016; Kanu *et al*. 2016; Nikkilä *et al*. 2017) almost certainly have DNA replication stress as a contributor, and it seems likely that the specific contexts driving replication stress will be important determinants of APOBEC3 mutation patterns.

Perhaps the most surprising finding yielded by analysis of APOBEC3C mutations is the absence of DNA strand biases that are evident in mutations caused by APOBEC3A and APOBEC3B. Although the importance of transcription bubbles as APOBEC3 targets in human cells remains unresolved (Nordentoft *et al*. 2014; Kazanov *et al*. 2015; Chervova *et al*. 2021; McCann *et al*. 2023), it is clear that in the humanized yeast models for APOBEC3A, APOBEC3B, and APOBEC3C there are different biases to the 5’ regions of tRNA and pol II genes. APOBEC3A and 3C show similar biases to tRNA and pol II genes, whereas APOBEC3B lacks a bias to pol II genes and causes a much smaller proportion of deaminations at tRNA genes (Figure 5c and f). Both APOBEC3A and APOBEC3B prefer to deaminate the non-transcribed strand, yet APOBEC3C shows no bias towards the non-transcribed strand (Figure 5d and g). For DNA replication strands, APOBEC3A and APOBEC3B show asymmetry with preferences for the leading strand which APOBEC3C lacks (Figure 6e and f). In the humanized yeast models ssDNA at DNA replication forks or transcription bubbles is expected to be present in an identical fashion and any human cell specific modulators of APOBEC3 function are expected to be absent. Thus, the data indicate that the simple presence of a ssDNA target is not sufficient for deamination by APOBEC3 and suggest that intrinsic features of each APOBEC3 enzyme will continue to be important keys to understanding the specificities of APOBEC3 enzymes in genome mutagenesis.

## Acknowledgements

I thank Elçin Ünal for the pUB1306 plasmid, Damiano Fantini for assistance with the MutSignatures package, Tajinder Ubhi, Brandon Ho, Raphael Loll-Krippleber, and Atina Coté for coding advice, Linda Chelico for helpful discussions, Jiongwen Ou for performing the immunoblots, and Tajinder Ubhi and members of the Brown lab for careful reading of the manuscript and helpful discussions. I am grateful to work on the lands of the Mississaugas of the Credit, the Anishnaabeg, the Haudenosaunee and the Wendat peoples, land that is now home to many diverse First Nations, Inuit, and Métis peoples.

