## Supplemental Figures for "The cytidine deaminase APOBEC3C has unique sequence and genome feature preferences"

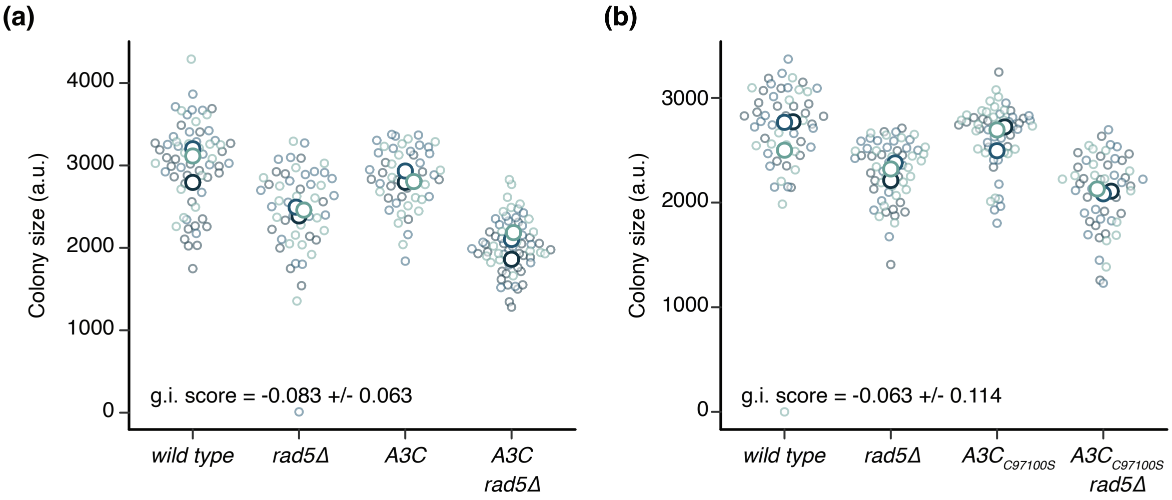


**Fig. S1.** Genetic interactions between *APOBEC3C* and *rad5Δ*. a) Colony size in arbitrary units (a.u.) was measured for the indicated strains following dissection of tetrads from *rad5Δ* x *APOBEC3C* crosses. Small circles show individual colony sizes (13 to 27 per genotype per replicate) and the large circles show the mean colony size for each of the 3 replicates. The genetic interaction (g.i.) score is indicated. b) Colony size in arbitrary units (a.u.) was measured for the indicated strains following dissection of tetrads from *rad5Δ* x *APOBEC3C$_{C97100S}$* crosses. Small circles show individual colony sizes (15 to 23 per genotype per replicate) and the large circles show the mean colony size for each of the 3 replicates. The genetic interaction (g.i.) score is indicated.

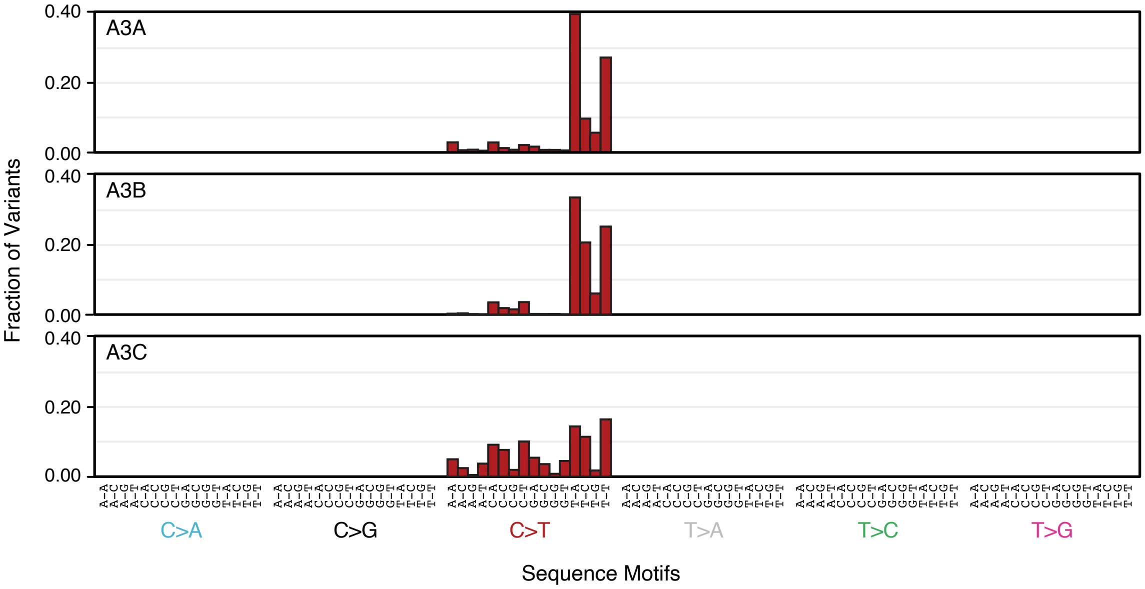


**Fig. S2.** APOBEC3 mutational signatures. The complete 96 tri-nucleotide mutational signatures are plotted for APOBEC3A, 3B, and 3C.

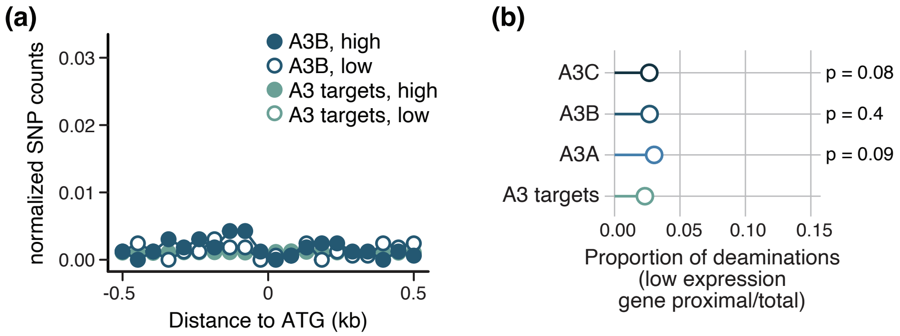


**Fig. S3.** Cytidine deaminations near the highest and lowest expressed genes. a) The proportion of SNVs in 50 bp bins surrounding the highest 5% (closed circles) and the lowest 5% of expressed genes is plotted for APOBEC3B. Putative APOBEC3 targets (TC) are plotted as a control. b) The proportion of deaminations +/- 500 bp of the lowest expressed genes is plotted for APOBEC3C, 3B, and 3A. Putative APOBEC3 targets (TC) are plotted as a control. Statistical support for enrichments proximal to highly expressed genes relative to APOBEC3 targets was assessed with chi-squared tests.

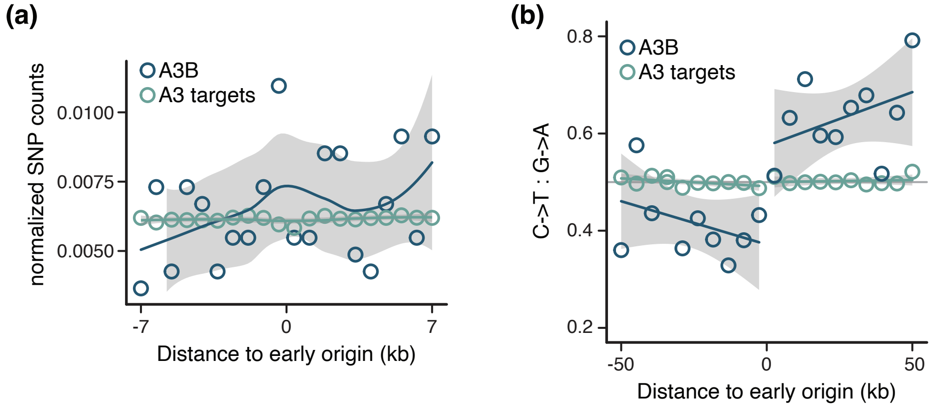


**Fig. S4.** APOBEC3B deaminations near early-firing DNA replication origins. a) The proportion of SNVs in 70 bp bins surrounding early-firing DNA replication origins is plotted for APOBEC3B. Putative APOBEC3 targets (TC) are plotted as a control. The solid line is the loess smoothed conditional mean with the shaded regions showing the standard error with a 0.95 confidence interval. b) The proportion of C->T SNVs in 5 kbp bins surrounding early-firing DNA replication origins is plotted for APOBEC3B. Putative APOBEC3 targets (TC) are plotted as a control. The solid line is the loess smoothed conditional mean with the shaded regions showing the standard error with a 0.95 confidence interval.
